# Lateralized alpha activity and slow potentials shifts over visual cortex track the time course of both endogenous and exogenous orienting of attention

**DOI:** 10.1101/2019.12.12.874818

**Authors:** Jonathan M. Keefe, Viola S. Störmer

**Author notes:** **Corresponding Author:** Jonathan Keefe.

## Abstract

Spatial attention can be oriented endogenously, based on current task goals, or exogenously, triggered by salient events in the environment. Based upon literature demonstrating differences in the time course and neural substrates of each type of orienting, these two attention systems are often treated as fundamentally distinct. However, recent studies suggest that rhythmic neural activity in the alpha band (8-13Hz) and slow waves in the event-related potential (ERP) may emerge over parietal-occipital cortex following both endogenous and exogenous attention cues. To assess whether these neural changes index common processes of spatial attention, we conducted two within-subject experiments varying the two main dimensions over which endogenous and exogenous attention tasks typically differ: cue informativity (spatially predictive vs. non-predictive) and cue format (centrally vs. peripherally presented). This task design allowed us to tease apart neural changes related to top-down goals and those driven by the reflexive orienting of spatial attention, as well as examine their interactions in a novel hybrid cross-modal attention task. Our data demonstrate that both central and peripheral cues elicit lateralized ERPs over parietal-occipital cortex, though at different points in time, consistent with these ERPs reflecting the orienting of spatial attention. Lateralized alpha activity was also present across all tasks, emerging rapidly for peripheral cues and sustaining only for spatially informative cues. Overall, these data indicate that distinct slow-wave ERPs index the spatial orienting of endogenous and exogenous attention, while lateralized alpha activity represents a common signature of visual enhancement in anticipation of potential targets across both types of attention.

## Introduction

Selective spatial attention can be deployed endogenously (i.e., voluntarily), following the goals and intentions of an observer, or exogenously (i.e., involuntarily), through capture by a sudden event in the environment such as a bright flash or a salient sound (Reynolds & Chelazzi, 2004; Wright & Ward, 2008). Decades of research have shown that endogenous and exogenous attention result in behavioral benefits at the attended location, reflected in higher accuracy and faster response times in discrimination or detection tasks (Posner, 1980; Posner & Cohen, 1984; for a review, see Carrasco, 2011). However, these behavioral effects typically arise at different timescales, with endogenous attentional benefits emerging slowly and sustaining for an extended time and exogenous attentional benefits emerging quickly but disappearing shortly after (with the possibility of behavioral costs thereafter, i.e., inhibition of return; Müller & Rabbit, 1989; Nakayama & Mackeben, 1989; Klein, 2000). Due to these differences in time course and origin (internal goals vs. external events), it is often assumed that these two modes of attention are fundamentally different.

This dissociation is supported by other evidence demonstrating that different neural substrates are involved in each type of attention, with partially separable fronto-parietal networks being responsible for the exogenous and endogenous orienting of attention (Hahn et al., 2006; Chica, Bartolomeo, & Lupiáñez, 2013; Corbetta & Shulman, 2002) and unique changes in fronto-parietal connectivity emerging following each type of orienting (Bowling, Friston, & Hopfinger, 2019). There is also evidence that the changes in visual-cortical activity resulting from the orienting of attention differ between these two systems. For example, human electrophysiological studies suggest that early visual-cortical processing is affected differently depending upon how attention is deployed, with exogenous attention affecting early visual processing of targets more strongly (P1 component of the visually-evoked potential) and endogenous attention affecting later processing more strongly (indexed by the N1 and P3 components; Hopfinger & West, 2006). Separate work recording single-unit activity in macaque MT demonstrates a similar pattern of earlier changes in firing rate following the onset of attention-grabbing (i.e. exogenous) cues and later changes following only cues that are relevant to the endogenous orienting of attention – further suggesting that separable mechanisms underlie each type of attention (Busse, Katzner, & Treue, 2008). These findings corroborate other research demonstrating that only informative cues elicit gamma-frequency EEG activity (Landau et al., 2007), which has been linked broadly to cognitive processes including attention (Tallon-Baudry, 2009). Altogether, findings like these have been taken as evidence that exogenous and endogenous attention represent two attention systems that affect sensory processing in different ways (for a review, see Chica et al., 2013).

However, other recent studies point to some commonalities in how the endogenous and exogenous orienting of attention affect neural processing in early sensory areas even prior to the onset of a target stimulus. Such early-arising similarities seem surprising given that both types of attention are initiated differently (salient bottom-up signals vs. endogenous top-down goals), and would imply that the behavioral benefits are supported by the same neural mechanisms. There are two particularly strong markers of these cue-triggered changes in neural activity observed in the electroencephalogram (EEG). During endogenous attention, the most commonly observed index is lateralized changes in the occipital alpha rhythm, an 8-13 Hz oscillation that tends to decrease over occipital areas contralateral to an attended location while increasing at ipsilateral sites in response to an attention cue. This lateralized decrease in alpha has been interpreted as representing the anticipatory biasing of visual-cortical activity in preparation of an impending target (Worden et al., 2000; Kelly et al., 2006; Green & McDonald, 2010). These changes in alpha activity have been shown to occur in endogenous cueing tasks around the same time as a slow-wave in the event-related potential (ERP), termed the Late-Directing Attention Positivity (LDAP), which has been interpreted as either reflecting pre-target biasing of visual activity (similar to the alpha changes) or the orienting of spatial attention itself (Harter et al., 1989; Hopf & Mangun, 2000; Eimer, Van Velzen, & Driver, 2002; Green & McDonald, 2006; Störmer, Green, & McDonald, 2009). Both of these lateralized changes over occipital and parietal-occipital cortex typically emerge relatively late after the onset of an attention cue (~ 500 to 700 ms later), in line with the slow time course of endogenous attention. Consequently, changes in alpha activity and the slow potential shift over parietal-occipital areas have both been interpreted as unique signatures of the endogenous orienting of attention. However, a different set of studies provides initial evidence that this might not be the case. These studies show that peripheral, salient sounds sometimes used to induce exogenous shifts of spatial attention can modulate the occipital alpha rhythm (Störmer et al., 2016; Feng et al., 2017) and trigger slow positive deflections in the ERP (McDonald et al., 2013; Feng et al., 2014; Störmer, 2019).

However, to date it is unclear how these alpha changes and slow-wave ERPs found across different studies relate to each other, and whether they represent common mechanisms associated with both types of attention. Given that these neural effects have been studied in very different paradigms using diverse stimuli and separate participants, the inferences that can be made in this regard are limited. A direct comparison of how exogenous and endogenous attention influence neural processing prior to the onset of a target is therefore lacking. Furthermore, and of particular interest to the current study, it is unclear how these changes in neural processing triggered by different cue types interact when both modes of attention are engaged.

To fill this gap, we conducted two within-subject experiments and varied the two main dimensions over which endogenous and exogenous attention tasks typically differ – cue informativity (spatially predictive vs. non-predictive) and cue format (centrally vs. peripherally presented) – while holding all other parameters constant. Across all tasks, we used auditory cues to orient spatial attention to avoid any contamination of visually evoked responses elicited by a visual cue. This allowed us to isolate neural activity related to the effects of the spatial orienting of attention. Importantly, the design also included a hybrid attention task with peripheral cues that were spatially predictive, combining aspects of both exogenous and endogenous attention. Our analysis focused on the temporal dynamics of slow-wave ERPs and lateralized alpha to disentangle processes related to the shifting of spatial attention, and resulting changes in visual-cortical excitability, triggered by salient bottom-up cues and top-down attentional goals.

With regards to the ERPs, we were particularly interested in whether the slow positive deflections elicited by endogenous (i.e., LDAP) and exogenous (i.e., ACOP) cues reflect shifts of spatial attention, sensory enhancement in visual areas, or a combination of the two. If these ERPs reflect the shifting of spatial attention to a new location, we would expect them to occur earlier for peripheral relative to central cues. Importantly, in the hybrid attention task, a positive deflection should only be present early (and not later), as attention would already have been shifted by the peripheral cue. Alternatively, if these slow positive deflections reflect pre-target biasing of visual-cortical activity, or a combination of shifting and pre-target biasing, we would expect them to be present continuously throughout the cue-target interval during endogenous and hybrid cueing tasks – in which the cues contain reliable information about the location of a future target.

With regards to alpha oscillations, we expected lateralized occipital alpha activity to emerge relatively early following exogenous cues and later following endogenous cues (Feng et al., 2017; Kelly et al., 2006). The main question of interest was whether these alpha changes represent a common process of biasing visual-cortical activity across both types of attention. The hybrid attention task (peripheral, informative cues) allowed us to test this directly. Specifically, if alpha indexes the same biasing process, then we would expect occipital-parietal lateralized alpha activity to be continuously present through both early and late time windows. Alternatively, if there is a discontinuity in this alpha activity or a large change in topography across time (e.g. only occipital or only parietal), then this would suggest that alpha activity indexes separate biasing processes following each type of attentional orienting. In addition, comparing the magnitude and topography of lateralized alpha across exogenous and hybrid attention tasks allowed us to examine the sensitivity of early alpha changes to top-down task goals (i.e., spatial informativity of the cue).

To anticipate our results, we find that lateralized alpha represents a common index of visual-cortical biasing across both modes of attention, but that the slow-wave ERPs -- ACOP and LDAP-- represent partially differentiable attentional orienting processes.

## Method

### Participants

Sixteen participants were included in the final sample of Experiment 1 (11 female; mean age of 21.9 years) and another 16 participants were included in the final sample of Experiment 2 (12 female; mean age of 21.7 years). Each of the subjects participated in only one of the experiments. For Experiment 1, data from three participants were excluded due to performance at or below chance level across all conditions (<~50% accuracy). An additional two participants did not complete the EEG task due to an inability to suppress saccades to the cue and/or target in initial practice tasks. For Experiment 2, data from three participants were excluded due to excessive artifacts in the EEG (affecting > 33% of trials). Data from an additional participant were excluded due to inability to perform the task, as the participant reported seeing two targets of orthogonal orientation at the same location – leading them to report guessing the orientation of the target on every trial.

All participants gave informed written consent as approved by the Human Research Protections Program of the University of California, San Diego and were paid for their time ($10/hour) or received course credit. All participants reported having normal or corrected-to-normal vision and normal hearing. Sample sizes were chosen a priori based upon a number of other studies utilizing similar cross-modal attentional cueing paradigms (McDonald, Teder-Salejarvi, & Hillyard, 2000; Green & McDonald, 2006; Störmer, McDonald, & Hillyard, 2009; McDonald et al., 2013; Feng et al., 2014).

### Stimuli and Apparatus

Participants were seated approximately 45 cm in front of a 27” monitor in a sound-attenuated, electrically shielded booth. Stimuli were presented on the screen via the Psychophysics Toolbox in MATLAB (Brainard, 1997; Pelli, 1997). A small black fixation dot (0.2° x 0.2° of visual angle) was always present in the center of the screen, which was otherwise uniformly gray (RGB: 127, 127, 127). A black circle (0.4° x 0.4°) appeared around the fixation dot at the start of each trial to indicate to the participant that the trial had begun. We ran three different tasks across the two experiments that differed only in the type of cues that were presented. In the hybrid (informative peripheral cues; Experiments 1 and 2) and exogenous (uninformative peripheral cues; Experiment 1) attention tasks, the cues were ~83 ms pink noise bursts (0.5–15 kHz, 78 dB SPL) played from external speakers mounted on either side of the computer monitor. The auditory stimuli were played in stereo and their amplitude was adjusted to give the impression that the sounds were emanating from the possible target locations on the screen. In the endogenous attention task (informative central cues; Experiment 2), the attention cue was either an upward frequency sweep ranging from 750 Hz to 1,000 Hz or a downward frequency sweep from 1,250 Hz to 1,000 Hz, played from both speakers and perceived emanating from the center/entirety of the screen. Across all tasks, the target was a Gabor patch with a spatial frequency of 1.3 cycles/degree, turned either −45° or 45° from vertical. The contrast of the Gabor patch was determined for each participant in a calibration task prior to the main experiment (see below). The target was presented in one of two peripheral locations indicated by a black circle with a diameter of ~9° visual angle, centered ~28° of visual angle to the left and right of fixation. Each target was followed by a visual noise mask of the same size.

### Experiment 1 Procedures

In Experiment 1 we compared whether and how the changes in parietal-occipital activity elicited by a peripheral and spatially informative cue differ relative to a peripheral and spatially uninformative cue usually used in exogenous attention tasks. In other words, these cues were physically identical but differed as to whether they indicated where the target was likely to appear. This allowed us to isolate the rapid effects of exogenous attention upon visual-cortical processing, triggered by reflexive shifts of attention to salient and peripheral cues, from the later effects of endogenous attention, which are triggered only by cues that carry temporal/spatial information about a target. All participants performed two cross-modal attention tasks, outlined in Figure 1A: the hybrid attention task and the exogenous attention task.

**Figure 1.**
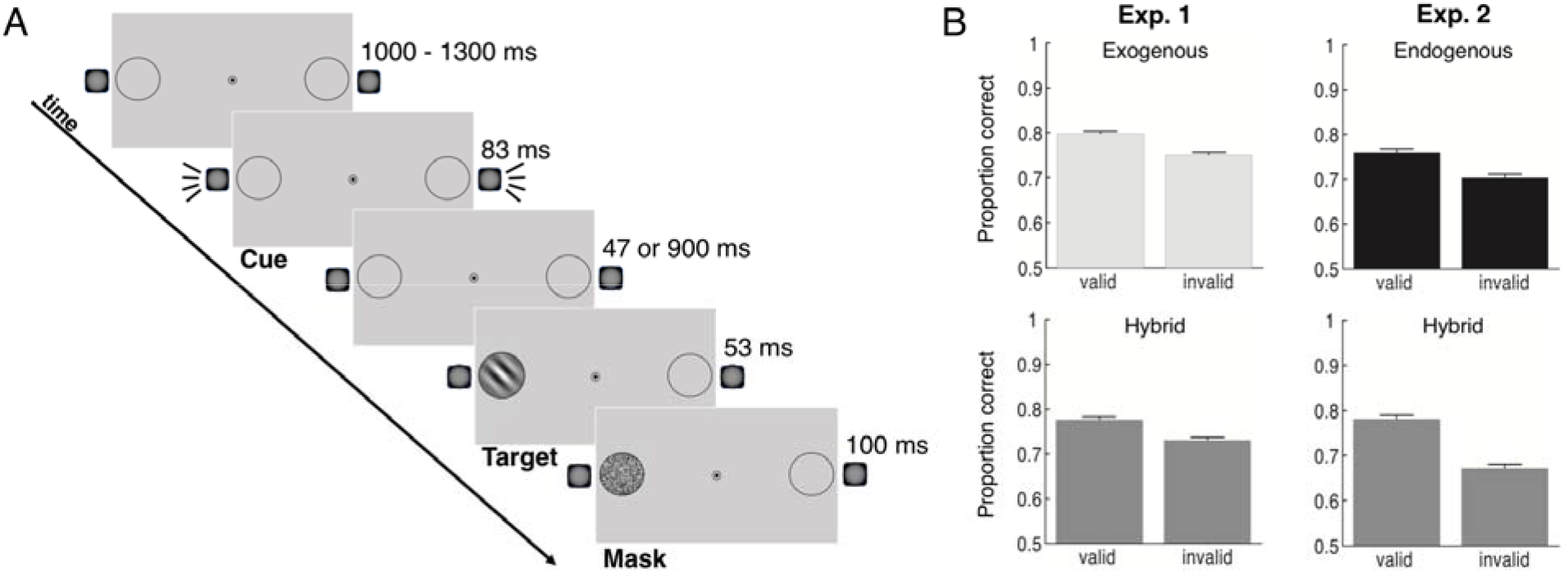
General task design and performance. (A) Participants discriminated the direction of rotation (clockwise or counterclockwise) of a masked Gabor patch target. Prior to the appearance of the target, participants were presented with an auditory cue that was either presented randomly 130 ms prior to the target (50% valid; Exogenous task), or informative (80% valid) as to the future target’s location following a ~1000 ms SOA (Endogenous and Hybrid tasks). This sound was a lateral pink noise burst in the hybrid and exogenous attention tasks, and an up-sweeping or down-sweeping tone in the endogenous attention task. (B) Target discrimination accuracy, plotted as a function of cue validity for each of the tasks in Experiment 1 and Experiment 2, clearly reveals a benefit in accuracy at the cued vs. uncued location across all tasks. Error bars represent ± 1 standard error of the mean.

In the hybrid attention task, participants were asked to keep their eyes on the central fixation dot throughout each experimental block. A black circle appeared around the central fixation dot at the beginning of each trial, indicating to the participants that the trial had begun. Following the onset of this circle at a variable stimulus onset asynchrony (SOA) of 1,000 – 1,300 ms, an 83-ms auditory attention cue was presented that indicated the location of a subsequent target with 80% validity (Posner, 1980). Participants were informed about the relationship between cue and target location and were thus instructed to covertly shift their attention to the cued side in anticipation of the target. After a cue-target SOA of ~980 ms, a Gabor patch target oriented 45° either clockwise or counterclockwise from vertical was presented at one of the two peripheral locations for ~53 ms and was followed immediately by a visual noise mask for 100 ms. The noise mask always appeared at the location of the target to eliminate uncertainty about the location at which the target appeared. Following the noise mask at an ISI of 300 ms, the black circle surrounding the central fixation dot turned white, prompting a response from the participant as to which direction the target was oriented. Participants made this report using the “m” (clockwise) and “n” (counterclockwise) keys.

The exogenous attention task differed in three ways from the hybrid attention task. First, instead of being informative as to where a future target would appear, the cue of the exogenous attention task was presented randomly at the left or right side and did not carry any spatial information about the target. Consequently, participants were instructed to ignore the cue because it would not be informative to the task. Second, the cue-target stimulus-onset asynchrony (SOA) in the exogenous task was much shorter than in the hybrid attention task (130 ms vs. ~980 ms) in order to eliminate any effects of endogenous attention and maximize the effects of exogenous attention. Third, a target was only presented on a randomly selected 50% of trials in the exogenous attention task. This was done in order to separate the neural activity elicited by the uninformative peripheral cue and the target, which would otherwise overlap given the short SOA. This design allowed us to isolate neural activity elicited by the cue without the contamination of activity elicited by the visual target. Thus, the analysis of behavioral performance was performed only on trials in which a target appeared and the analysis of EEG activity was performed only on trials in which a target did not appear (i.e., cue-only trials). On the trials in which a target was not presented, participants were asked to keep their eyes on the central fixation dot and prepare for the next trial.

All trial types were randomly intermixed, but the task performed (exogenous vs. hybrid) was blocked and the order counterbalanced between participants, such that half of the participants started with the exogenous attention task and the remaining half started with the hybrid attention task. The hybrid attention task consisted of 7 consecutive blocks of 48 trials each, whereas the exogenous attention task consisted of 7 consecutive blocks of 96 trials each in order to collect a comparable number of ERP epochs and behavioral trials for the analysis across both tasks. Note that each experimental block took approximately the same amount of time since the trial time was much shorter in the exogenous attention task. Prior to the experimental tasks, task difficulty was adjusted for each participant using a thresholding procedure that varied the contrast of the Gabor patch target to achieve about 75% accuracy (i.e., QUEST; Watson & Pelli, 1983). In this thresholding task, participants discriminated the direction of the 45°-oriented Gabor patch in the absence of any sounds. Each participant performed 72 trials of the thresholding task and the individual contrast thresholds were used for the main experiment. Participants performed 32 practice trials prior to each task.

### Experiment 2 Procedures

In Experiment 2, we compared whether and how the changes in parietal-occipital activity elicited by a peripheral and spatially informative cue differ relative to a central and spatially informative cue usually used in endogenous attention tasks. In other words, these cues conveyed the same information regarding where the target was likely to appear but differed in their physical properties such that only one cue conveyed spatial information itself. This allowed us to isolate the effects of endogenous attention upon parietal-occipital processing, triggered by voluntary shifts of attention to cues that carry temporal/spatial information about a target, from the earlier effects of exogenous attention, which are triggered by salient and peripheral cues that capture attention. All participants performed both the endogenous attention task and the hybrid attention task.

The hybrid attention task was identical to the task described in Exp. 1 procedures. The hybrid and endogenous attention tasks differed only in the type of auditory cue presented. In the hybrid attention task, the cue was a pink noise burst presented at either the left or right speaker and the location of the cue indicated where the target was likely to appear. In the endogenous attention task, participants were presented with a centrally presented up-sweeping or down-sweeping tone on each trial. The direction of the frequency sweep of that tone (up or down) indicated where the target was likely to appear on that trial (left or right; cf., Störmer, Green, McDonald, 2009). The sweep-direction-to-location mapping of this cue was counterbalanced across participants, such that the up-sweeping tone indicated that the target was likely to appear on the right side for half of the participants and the left side for the remaining half of participants. These different cue formats were chosen in order to dissociate the purported endogenous and exogenous components of attention; while the peripheral noise burst involved aspects of both exogenous and endogenous spatial attention tasks (i.e., peripherally presented and spatially predictive), the centrally presented sweeping tone involved aspects of only the traditional endogenous spatial attention tasks (i.eg., symbolic central cues that are spatially predictive).

All trial types were randomly intermixed, but the task performed (hybrid vs. endogenous) was blocked and the order counterbalanced between participants. Prior to performing either of the tasks, task difficulty was adjusted for each participant using the thresholding procedure described in Exp. 1. Participants performed 7 consecutive blocks of 48 trials for each task, after completing 32 practice trials in each task. An additional short practice block (24 trials) was performed prior to the endogenous attention task in order to familiarize participants with the symbolic auditory cues. In this practice task, participants were presented the up-sweeping and down-sweeping tones and asked to report the side on which the cue indicated the target would be likely to appear, in the absence of any visual information on the screen.

### EEG Recording and Analysis

Electroencephalogram (EEG) was recorded continuously from 32 Ag/AgCl electrodes mounted in an elastic cap and amplified by an ActiCHamp amplifier (BrainProducts, GmbH). Electrodes were arranged according to the 10-20 system. The horizontal electrooculogram (HEOG) was recorded from two additional electrodes placed on the external ocular canthi which were grounded with an electrode placed on the neck of the participant. The vertical electrooculogram was measured at electrodes FP1 or FP2, located above the left and right eye, respectively. All scalp electrodes were referenced to the right mastoid online and were digitized at 500 Hz.

Continuous EEG data were filtered with a bandpass (butterworth filter) of 0.01-112.5Hz offline. Data were epoched from −1,000 ms to +2,000 ms with respect to the onset of the auditory cue. Trials contaminated with blinks, eye movements, or muscle movements were removed from the analysis. Artifacts were detected in the time window −800 to 1,100ms in two steps. First, we used automated procedures implemented in ERPLAB (Lopez-Calderon & Luck, 2014; peak-to-peak for blinks, and a step function to detect horizontal eye movements at the HEOG channel). Second, for each participant, each epoch was visually inspected to check the automated procedure and the trials chosen for rejection were updated (cf., Störmer, Alvarez, & Cavanagh, 2014). To ensure that eye movements were fully removed using this artifact rejection method, we checked for any differences from zero in the HEOG channel separately for leftward and rightward cues. Specifically, we performed the same statistical analysis as for the time course analysis of ERPs and alpha (checking if 4 or more consecutive time bins of 50ms each were significantly different from zero; see *Statistical Analysis*), and we found no reliable drifts from baseline in any of the conditions. Artifact-free data was digitally re-referenced to the left mastoid. For the endogenous and hybrid attention tasks, all trials were included in the EEG analysis. For the exogenous attention task, only trials with no target stimuli were included to avoid overlap of the target-elicited neural activity with the cue-elicited neural activity.

ERPs elicited by the left and right noise bursts were averaged separately and were then collapsed across sound position (left, right) and hemisphere of recording (left, right) to obtain waveforms recorded ipsilaterally and contralaterally relative to the sound. The ERPs elicited by the central cues (up- and down-sweeping tones) were averaged separately for attend-left and attend-right conditions and then also collapsed across hemisphere and hemifield. ERPs were low-pass filtered (half-amplitude cutoff at 30 Hz; slope of 12dB/octave) to remove high-frequency noise. Mean amplitudes for each participant and condition were measured with respect to a 200 ms prestimulus period (−200 to 0 ms from cue onset), and mean amplitudes were statistically compared using both repeated-measures Analyses of Variance (ANOVAs) and paired t-tests (contralateral vs. ipsilateral to attended location). Our analysis was focused on two ERP components that have previously been associated with exogenous and endogenous spatial attention. In particular, we examined the Auditory-Evoked Contralateral Occipital Positivity (ACOP) as an index of exogenous attention (McDonald et al., 2013), and the Late-Directing Attention Positivity (LDAP) as a signature of endogenous attention (Harter et al., 1989; Eimer et al., 2002; Green & McDonald, 2006). The exact time windows and electrode sites for each ERP analysis were chosen a priori based on previous research and matched across all analyses. Both ERP components were measured at the same four parietal-occipital electrode sites (PO7/PO8/P7/P8). These electrodes were chosen because they are typically used in ACOP paradigms (McDonald et al., 2013; Feng et al, 2017). Though prior LDAP studies have focused mainly on analyzing activity at PO7/8 (Green & McDonald, 2008; 2010), we included P7/8 to keep analyses consistent across the components. The ACOP was measured between 260-360 ms (McDonald et al., 2013), while the LDAP was measured between 500 - 800 ms (Green & McDonald, 2006). Additional pairwise comparisons (contralateral vs. ipsilateral) were performed on successive 50 ms sections of the ERP in order to better characterize the time course of these positive deflections in each task (cf., McDonald & Green, 2008; Störmer et al., 2009).

For the time frequency analysis, scalp channels were analyzed via complex Morlet wavelets before averaging, following the methods of Lakatos et al. (2004) and Torrence and Compo (1998). Spectral amplitudes were calculated via four-cycle wavelets at 60 different frequencies increasing linearly from 2 to 40 Hz separately for each electrode, time point (every 2 ms), attention condition (left, right), and participant. Spectral amplitudes were then averaged across trials separately for each condition and participant, and a mean baseline of –350 to –150 ms from cue onset was subtracted from each time point for each frequency separately (Pitts, Padwal, Fennelly, Martínez, & Hillyard, 2014; Störmer et al., 2016). Mean spectral amplitudes elicited by the left and right noise bursts (exogenous and hybrid attention tasks) and left- and right-directing central tones (endogenous attention task) were then collapsed across cued location (left, right) and lateral position of the electrode (left, right) to reveal attention-induced modulations ipsilateral and contralateral to the cued location. The statistical analysis was focused on alpha-band amplitude modulations over the range of 8 – 13 Hz at parietal-occipital electrode sites (PO7/PO8/P7/P8) and during the same time intervals as the ACOP (260 – 360 ms) and LDAP (500 – 800 ms) components. Replicating the ERP analysis, pairwise comparisons were performed on successive 50 ms sections of the average alpha-band amplitude values (i.e., average amplitude of oscillatory activity across 8-13 Hz) of the ipsilateral and contralateral hemispheres in each task. Data processing was carried out using EEGLAB (Delorme & Makeig, 2004) and ERPLAB (Lopez-Calderon & Luck, 2014) toolboxes and custom-written scripts in MATLAB (The MathWorks, Natick, MA).

### Topographical maps

To illustrate the scalp distribution of the different ERP and time-frequency measures, we created topographical maps using spline interpolation of the voltage differences between the contralateral and ipsilateral hemispheres for each of the time windows of interest. Specifically, the contralateral-minus-ipsilateral ERPs and alpha activity difference were calculated for homologous left and right electrode sites (e.g., PO7 and PO8), with the values at midline electrode sites (e.g., POz) set to zero (Störmer et al., 2009). These difference voltage topographies were projected to the right side of the head.

### Statistical Analyses

Behavior was analyzed by comparing accuracy (% correct) in the Gabor discrimination task separately for when the Gabor patch appeared at the cued location (valid trials) vs. at the uncued location (invalid trials). Behavioral and EEG data were statistically analyzed using paired t-tests and repeated-measures ANOVAs (alpha = 0.05) using MATLAB (The MathWorks, Natick, MA). In order to control for spurious results in the time window analyses of the EEG data, a statistical difference between the activity of each hemisphere in a time window was only considered reliable if it was significant and was a part of a cluster of four or more significant time windows (i.e., there were 4 or more consecutive time windows with *p* < .05; Luck, 2014).

## Results

### Exp. 1 Behavior

As shown in Figure 1B, accuracy was higher following valid vs. invalid cues in both the exogenous and hybrid attention tasks of Experiment 1. In order to confirm the presence of this behavioral cueing benefit in each task, a two-way repeated-measures ANOVA with factors of cue validity (valid or invalid) and task (endogenous or hybrid) was performed. This analysis revealed a significant main effect of cue validity, *F*(1, 15) = 33.42, *p* < 0.001, η^2^ = 0.09, confirming that the higher accuracy following valid than invalid cues was reliable. There was no main effect of task, *F*(1, 15) = 1.38, *p* = 0.26, η^2^ = 0.02, nor an interaction between cue validity and task, *F*(1, 15) = 0.00, *p* = 0.95, η^2^ < 0.001, indicating that neither overall task performance nor the magnitude of the observed behavioral cueing benefits differed between tasks. Follow-up paired t-tests confirmed that accuracy was higher following valid than invalid cues in both the exogenous, *t*(15) = 3.87, *p* = 0.002, *d* = 0.97, and hybrid attention tasks, *t*(15) = 3.24, *p* = 0.006, *d* = 0.81.

In order to confirm that these differences in accuracy were not the result of a speed-accuracy trade-off, we also analyzed reaction times (i.e. RTs) to the target. Numerically, we found that RT in the exogenous task was roughly equivalent following valid cues (*M* = 1279 ms, *sd* = 997 ms) and invalid cues (*M* = 1291 ms, *sd* = 995 ms); and that RT in the hybrid task was faster following valid cues (*M* = 920 ms, *sd* = 543 ms) than invalid cues (*M* = 1029 ms, *sd* = 549 ms). In order to test whether this pattern held statistically, we performed a two-way repeated-measures ANOVA with factors of cue validity (valid or invalid) and task (endogenous or hybrid) on this RT data. This analysis revealed a significant main effect of cue validity, *F*(1, 15) = 7.57, *p* = 0.01, η^2^ = 0.001, but there was no main effect of task, *F*(1, 15) = 2.9, *p* = 0.11, η^2^ = 0.04, nor an interaction between cue validity and task, *F*(1, 15) = 1.35, *p* = 0.26, η^2^ < 0.001. Follow-up paired t-tests probing the main effect of cue validity confirmed that RT was not significantly different following valid and invalid cues in the exogenous task, *t*(15) = 0.21, *p* = 0.84, *d* = 0.05; and RT was significantly faster following valid than invalid cues in the hybrid attention task, *t*(15) = 2.63, *p* = 0.02, *d* = 0.66. These findings demonstrate that higher accuracy following the valid vs. invalid cues of each task in Experiment 1 cannot be explained by a trade-off between speed and accuracy.

### Exp. 1 Cue-elicited ERPs

Previous research has proposed that slow positive deflections in the ERP following informative central cues (the LDAP) and salient peripheral cues (the ACOP) may both represent either the orienting of spatial attention itself or the enhancement of visual-cortical processing prior to the onset of a target (Hillyard et al., 2016). If these positive deflections do in fact index a common process, we expect to observe positivities of similar topography in both the exogenous and hybrid tasks. However, the expected time course of these ERPs would vary based upon whether the ERPs commonly index the orienting of attention or the biasing of visual cortex itself. Specifically, if these slow potentials reflect the orienting of attention to a spatial location, we would expect to see a positivity of similar time course in response to each of the cues here – as attention should be exogenously shifted by the salient peripheral cues regardless of their informativity. Alternatively, if these positivities reflect the anticipatory biasing of visual-cortical activity, we would expect to observe only an early positivity in response to the uninformative cue of the exogenous task and both an early and late (or sustained) positivity in response to the informative cue of the hybrid attention task. Differences in spatial topography between these positivities or departures from the expected temporal patterns of these changes would argue against the interpretation of these positive ERPs as common indices of attentional orienting and/or visual-cortical biasing.

As shown in Figure 3A, the ERP waveforms were more positive over the hemisphere contralateral vs. ipsilateral with respect to the cued location during, and beyond, the ACOP time window (260 – 360 ms) of both the exogenous and hybrid attention tasks. A two-way repeated-measures ANOVA with factors of hemisphere (ipsilateral vs. contralateral) and task (exogenous vs. hybrid) was performed on the ERP waveforms during the ACOP time window. This analysis revealed a main effect of hemisphere, *F*(1, 15) = 20.88, *p* < 0.001, η^2^ = 0.07, indicating a significant difference between the amplitude of the ipsilateral and contralateral waveforms (i.e., ACOP). The magnitude of the ACOP was comparable across both tasks, as there was no significant main effect of task, *F*(1, 15) = 0.78, *p* = 0.39, η^2^ = 0.01, nor an interaction between hemisphere and task, *F*(1, 15) = 0.02, *p* = 0.90, η^2^ < 0.001.

**Figure 2.**
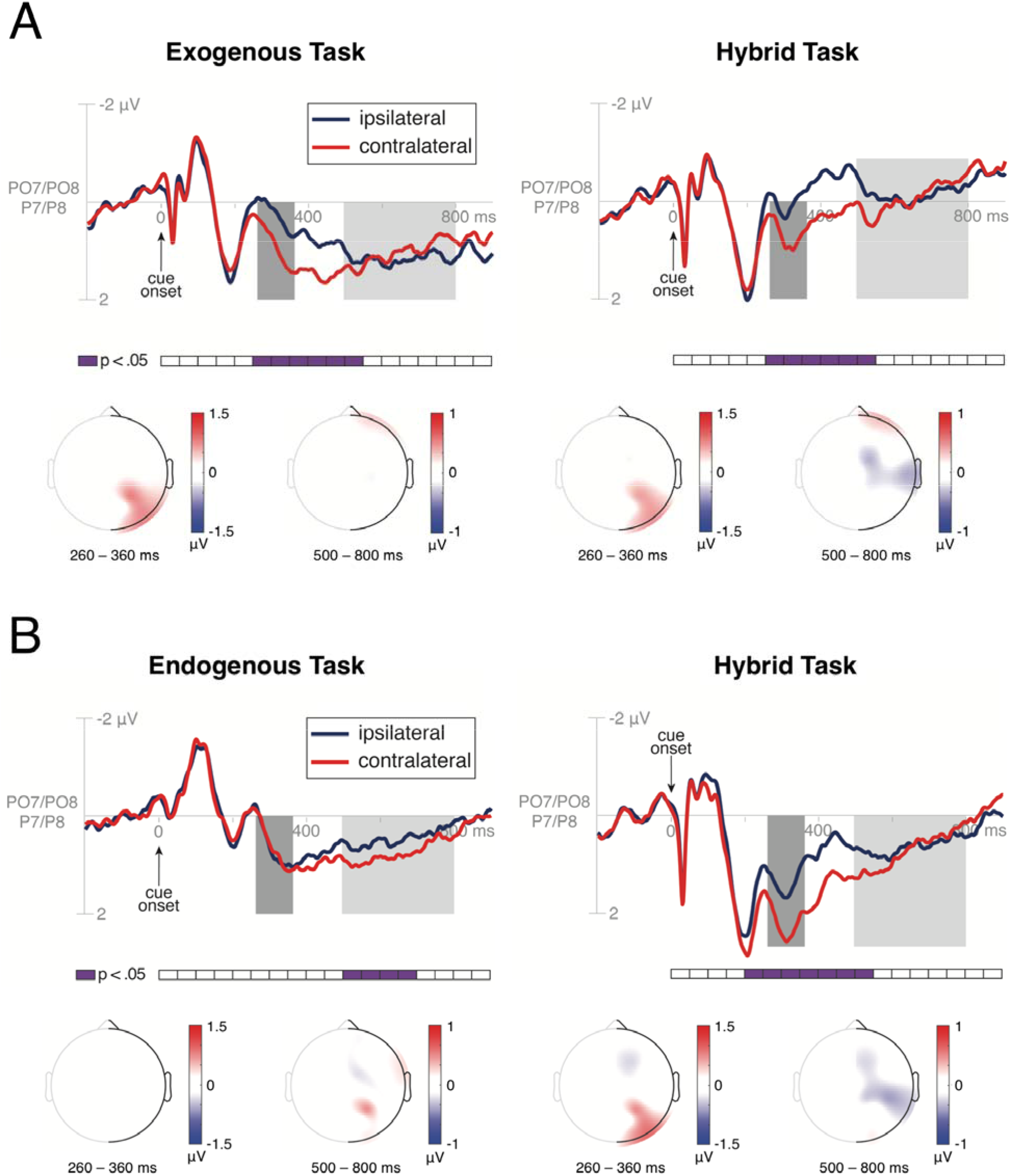
Grand-average ERP waveforms and topographies. ERPs at parietal-occipital scalp sites (PO7/PO8/P7/P8) were collapsed over left- and right-cue conditions and left and right hemispheres to obtain waveforms recorded ipsilaterally and contralaterally to the cued location. A priori defined ACOP and LDAP time windows are highlighted in dark gray and light gray, respectively. Statistically significant (p ≤ 0.05) differences between contralateral and ipsilateral waveforms are denoted in purple below the time axis. Topographical voltage maps show the contralateral-minus-ipsilateral ERP difference amplitudes, projected to the right side of the scalp during the ACOP and LDAP time windows. (A) A significant, early contralateral positivity (i.e. ACOP) was observed in response to the uninformative, peripheral cues of the exogenous attention task as well as the informative, peripheral cues of the hybrid attention task of Experiment 1. No LDAP was observed in the tasks containing peripheral sounds. (B) A significant late positivity (i.e. LDAP) contralateral to the cued location was observed in response to the symbolic, central cues of the endogenous attention task of Experiment 2. An earlier contralateral positivity (i.e. ACOP) was observed in response to the informative, peripheral cues of the hybrid attention task.

**Figure 3.**
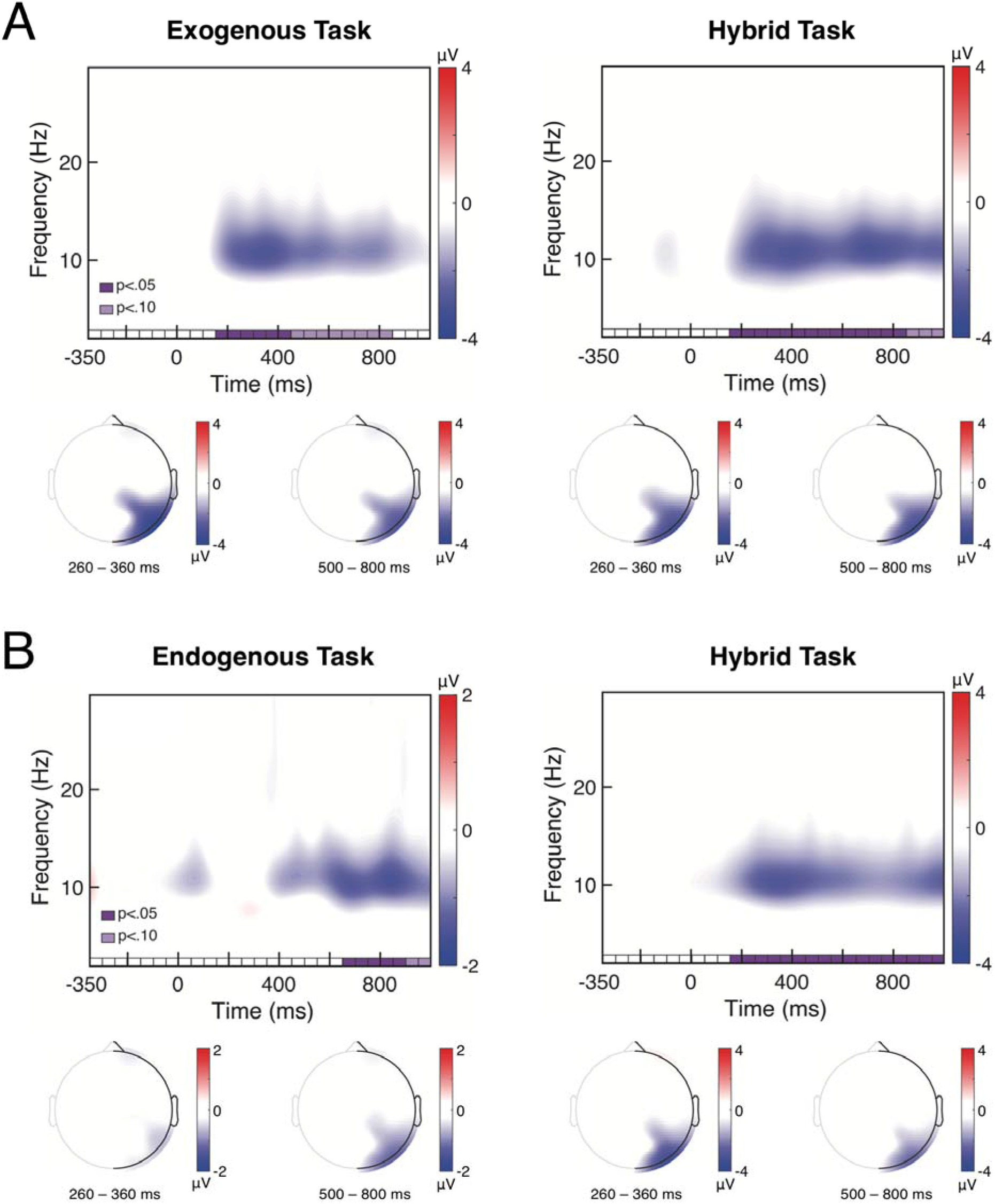
Grand-average time frequency plots of the contralateral-minus-ipsilateral activity over parietal-occipital scalp (PO7/PO8/P7/P8) shows clear lateralized alpha-band changes (8–13 Hz). Statistically significant (p ≤ 0.05; dark purple boxes) and near-significant (p ≤ 0.10; light purple boxes) differences between contralateral and ipsilateral alpha amplitude are denoted on the time axis. Topographical voltage maps show the contralateral-minus-ipsilateral alpha-band difference amplitudes, projected to the right side of the scalp, during the pre-defined ACOP and LDAP time windows. (A) Lateralized decreases in contralateral (relative to ipsilateral) alpha-band amplitude emerged rapidly following both the uninformative peripheral cues of the exogenous attention task and the informative, peripheral cues of the hybrid attention task of Experiment 1. (B) Contralateral decreases in alpha-band amplitude emerged following both the symbolic, central cues of the endogenous attention task and the informative, peripheral cues of the hybrid attention task of Experiment 2. Topographical maps show a clear contralateral occipital focus of the alpha changes in all conditions. Note the difference in the scale of the color bar in the endogenous task vs. other tasks.

Conversely, as can be seen in Figure 3A, a later contralateral vs. ipsilateral positivity (i.e., LDAP) was not readily evident in the ERP waveforms of either task. In order to test this statistically, a two-way repeated-measures ANOVA with factors of hemisphere (ipsilateral vs. contralateral) and task (exogenous vs. hybrid) was performed on the ERP waveform during the LDAP time window (500 – 800 ms). This analysis confirmed that there was no effect of hemisphere, *F*(1, 15) = 0.24, *p* = 0.63, η^2^ < 0.001, nor an interaction between hemisphere and task, *F*(1, 15) = 1.63, *p* = 0.22, η^2^ = 0.002, indicating that there was no hint of an LDAP. However, there was a significant main effect of task, *F*(1, 15) = 24.01, *p* < 0.001, η^2^ = 0.29, indicating a general difference in average ERP magnitude between the two tasks. Altogether, these results show that a reliable contralateral positivity of comparable magnitude emerged quickly after cue onset over occipital and parietal-occipital cortex (i.e., ACOP), regardless of whether the cue was spatially informative (hybrid task) or uninformative (exogenous task).

To examine the time course of these positivities in more detail, pairwise comparisons were performed on successive 50 ms sections of the ipsilateral and contralateral ERP waveforms of each task. These comparisons indicated that the ACOP stretched from 250 - 550 ms in both the exogenous attention task (all *p*s < 0.05) and the hybrid attention task (all *p*s < 0.04).

These data argue against an interpretation of the early and late positivities previously observed in exogenous and endogenous attention tasks as common indices of visual-cortical biasing in anticipation of a target, as there was no late or sustained response to the informative peripheral cues of the hybrid task. However, the emergence of an early positivity (i.e. ACOP) over occipital and parietal cortex in response to each of the peripheral cues suggests that the ACOP may represent a neural index of the orienting of attention and/or the initial biasing of visual-cortical processing. Notably, this orienting and/or biasing appears to occur regardless of the spatial informativity of the cue and is thus reflexive.

### Exp. 1 Cue-Elicited Alpha Oscillations

If lateralized changes in alpha activity are an index of a common process of visual-cortical facilitation across exogenous and endogenous attention, then we would expect to observe changes in alpha activity of similar topographies across each of the tasks of Experiment 1. Specifically, based upon prior literature, we might expect to observe a quick but relatively shortlived alpha change in response to the uninformative peripheral cues of the exogenous task and both an early and late (or sustained) alpha change in response to the informative peripheral cues of the hybrid task. Alternatively, if changes in alpha activity index processes unique to each type of attention, then we would expect to either not observe them in one of the tasks or find large differences in the topography of alpha modulations across tasks, which would suggest separate neural sources.

As demonstrated in the contralateral-minus-ipsilateral difference plots of Figure 2A, cues in both the exogenous and hybrid attention tasks elicited lateralized changes in alpha frequency amplitude over parietal-occipital cortex, such that there was a greater decrease in alpha amplitude contralateral relative to ipsilateral to the cued location. In order to probe the time course of this lateralized oscillatory alpha activity in each task, pairwise comparisons were performed on successive 50 ms sections of the average alpha-band amplitude values of the ipsilateral and contralateral hemispheres in each task. This analysis revealed significant lateralized alpha activity in the exogenous attention task from 150 – 450 ms (*p*s < 0.04), with marginally non-significant alpha activity stretching from 450 – 850 ms (*p*s < 0.10), and no reliable alpha activity thereafter. In contrast, significant lateralized alpha activity was present in the hybrid attention task from 150 – 850 ms (*p*s < 0.05), with marginally non-significant alpha activity from 850 – 1000 ms (*p*s < 0.07).

In order to compare the magnitude of this lateralized alpha activity across tasks, pairwise comparisons were performed on the alpha amplitude difference values (contralateral minus ipsilateral alpha amplitude) of each task in the a priori defined ACOP (260 – 360 ms) and LDAP (500 – 800 ms) time windows. These comparisons indicated that there was not a significant difference in the amplitude of lateralized alpha activity at the early time window, *t*(15) = 0.01, *p* = 0.99, *d* = 0.003, or at the late time window, *t*(15) = 1.49, *p* = 0.16, *d* = 0.37. In sum, these results show that lateralized alpha activity of similar magnitude and topography emerges rapidly (~150ms post cue) following peripheral auditory cues, regardless of their spatial informativity, but tends to decay earlier (~450ms) when the cue is not informative about the spatial location of a target relative to when it predicts the target location (~850ms).

### Exp. 2 Behavior

As shown in Figure 1B, accuracy was higher following valid vs. invalid cues in both the endogenous and hybrid attention task of Experiment 2. Following the analysis strategy of Exp. 1, a two-way repeated-measures ANOVA with factors of cue validity (valid vs. invalid) and task (endogenous vs. hybrid) was performed. There was no significant main effect of task, *F*(1, 15) = 0.15, *p* = 0.70, < 0.001, but there was a significant main effect of cue validity, *F*(1, 15) = 39.39, *p* < 0.001, η^2^ = 0.28, indicating that accuracy was significantly higher following valid relative to invalid cues. Interestingly, the magnitude of the observed behavioral benefits was greater in the hybrid attention task than in the endogenous attention task, as indicated by a significant interaction between cue validity and task, *F*(1, 15) = 5.91, *p* = 0.03, η^2^ = 0.03. Follow-up t-tests confirmed that accuracy was higher following valid than invalid cues in both the endogenous task, *t*(15) = 3.45, *p* = 0.004, *d* = 0.86, and hybrid attention task, *t*(15) = 6.12, *p* < 0.001, *d* = 1.53.

In order to confirm that these differences in accuracy were not the result of a speed-accuracy trade-off, we analyzed reaction times (i.e. RTs) to the target. Numerically, we found that RT in the endogenous task was faster following valid cues (*M* = 923 ms, *sd* = 737 ms) than invalid cues (*M* = 1089 ms, *sd* = 797 ms); and that RT in the hybrid task was faster following valid cues (*M* = 842 ms, *sd* = 450 ms) than invalid cues (*M* = 1022 ms, *sd* = 564 ms). In order to test whether this pattern held statistically, we performed a two-way repeated-measures ANOVA with factors of cue validity (valid or invalid) and task (endogenous or hybrid) on this RT data. This analysis revealed a significant main effect of cue validity, *F*(1, 15) = 10.07, *p* = 0.01, η^2^ = 0.02, but there was no main effect of task, *F*(1, 15) = 0.55, *p* = 0.47, η^2^ = 0.003, nor an interaction between cue validity and task, *F*(1, 15) = 0.04, *p* = 0.85, η^2^ < 0.001. Follow-up paired t-tests probing the main effect of cue validity confirmed that there was a marginally nonsignificant difference in RT following valid and invalid cues in the endogenous task, *t*(15) = 2.07, *p* = 0.06, *d* = 0.52; and that RT was significantly faster following valid than invalid cues in the hybrid attention task, *t*(15) = 3.88, *p* = 0.002, *d* = 0.97. These findings demonstrate that higher accuracy following the valid vs. invalid cues of each task in Experiment 2 cannot be explained by a trade-off between speed and accuracy.

### Exp. 2 Cue-elicited ERPs

Experiment 1 showed that the ACOP was elicited following peripheral informative and uninformative cues, indicating that the ACOP is robustly triggered by peripheral cues, regardless of their spatial predictability. However, it is unclear whether both the ACOP and LDAP both index the spatial shifting of attention to a new location, especially given that the LDAP is usually observed in endogenous tasks using central cues,. If this is the case, then we would expect to observe positivities of similar topography in the endogenous and hybrid tasks that differ only in their time course. Differences in spatial topography between these positivities or departures from the expected temporal patterns of these changes would argue against the interpretation of these positivities as common indices of attentional orienting.

As shown in Figure 3B, the ERP waveform contralateral to the cued location was more positive than the waveform ipsilateral to the cued location during, and beyond, the ACOP time window (260 – 360 ms) in the hybrid attention task. Conversely, this early positivity was not present in the endogenous attention task. To provide statistical support for these observations, a two-way repeated-measures ANOVA with factors of hemisphere (ipsilateral vs. contralateral) and task (endogenous vs. hybrid) was performed on the ERP waveform during the ACOP time window. This analysis revealed a significant main effect of hemisphere, *F*(1, 15) = 11.99, *p* = 0.004, η^2^ = 0.02, and task, *F*(1, 15) = 26.78, *p* = < 0.001, η^2^ = 0.14, as well as a significant interaction between hemisphere and task, *F*(1, 15) = 21.76, *p* < 0.001, η^2^ = 0.02, indicating that the main effects were driven by differences in the magnitude of the ACOP between tasks. Follow-up t-tests comparing the magnitude of the ipsilateral and contralateral ERP waveforms in each task revealed the presence of an ACOP in the hybrid attention task, *t*(15) = 4.52, *p* < 0.001, *d* = 1.13, but not the endogenous attention task, *t*(15) = 0.08, *p* = 0.94, *d* = 0.02.

Conversely, as can be seen in Figure 3B, a later contralateral vs. ipsilateral positivity (i.e., LDAP) was evident only in the ERP waveform of the endogenous attention task. In order to test for the presence of an LDAP in each task, a two-way repeated-measures ANOVA with factors of hemisphere (ipsilateral or contralateral) and task (endogenous or hybrid) was performed on the ERP waveform during the LDAP time window (500 – 800 ms). The analysis indicated that there was no significant main effect of task, *F*(1, 15) = 0.16, *p* = 0.70, η^2^ < 0.001, nor an interaction between task and hemisphere, *F*(1, 15) = 1.48, *p* = 0.24, η^2^ = 0.002. However, this analysis revealed a marginally non-significant main effect of hemisphere, *F*(1, 15) = 3.82, *p* = 0.07, η = 0.01. In order to probe this marginal effect further, follow-up t-tests were performed comparing the ipsilateral and contralateral ERP waveforms during the LDAP time window for each task. These comparisons indicated that there was no reliable LDAP in the hybrid attention task, *t*(15) = 0.60, *p* = 0.56, *d* = 0.15, but did indicate the presence of a significant LDAP in the goal-directed attention task, *t*(15) = 2.54, *p* = 0.02, *d* = 0.64. Altogether, these results indicate that a significant contralateral positivity emerged quickly following the informative, peripheral cue of the hybrid attention task (i.e., ACOP), and that an analogous – albeit smaller – contralateral positivity emerged on a later time frame (i.e., LDAP) following the informative, central cue of the endogenous attention task.

In order to examine the time course of each positivity in more detail, pairwise comparisons were performed on successive 50 ms sections of the ipsilateral and contralateral ERP waveforms of each task. These comparisons indicated the presence of a significant positivity from 200 – 550 ms in the hybrid attention task (i.e., ACOP; all *ps* < 0.02), and a significant positivity present from 500 – 700 ms (i.e., LDAP; all *ps* < 0.03) in the endogenous attention task.

Together, these data indicate that orienting attention following both peripheral and central informative cues results in a positivity in the cue-locked ERP. However, the time course of this positivity differs based upon the format of the cue – with a much earlier positive deflection emerging following peripheral cues and a relatively late positivity following central cues. Neither of these positivities sustained over the entire cue-target interval, further arguing against an account of the ACOP and LDAP as indices of the sustained biasing of visual-cortical processing. Additionally, these components appear to differ in their spatial topography, with the ACOP showing both occipital and parietal-occipital foci and the LDAP showing only a parietal focus. This difference in topography implies that each positivity indexes partially differentiable underlying processes (see Discussion).

### Exp. 2 Cue-Elicited Alpha Oscillations

In Experiment 1 we found that peripheral cues – regardless of their spatial informativity – elicited rapid changes in occipital alpha that sustained contiguously when the cue carried spatial information about the subsequent target, suggesting that lateralized alpha represents a common process underlying exogenous and endogenous attention. However, it is unclear from Experiment 1 whether the later changes in alpha activity observed in the hybrid task are similar to alpha changes elicited by central, symbolic cues typically used in endogenous cueing paradigms. If this is the case, then we would expect to observe late lateralized alpha activity of similar topography in response to informative peripheral and central symbolic cues of the hybrid and endogenous tasks, respectively.

As demonstrated in the contralateral-minus-ipsilateral difference plots of Figure 2B, both the endogenous and hybrid attention tasks elicited lateralized changes in alpha frequency amplitude, such that there was a greater decrease in alpha amplitude over the hemisphere contralateral relative to ipsilateral with respect to the cued location. First, in order to probe the time course of this lateralized alpha oscillatory activity in each task, pairwise comparisons were performed on successive 50 ms sections of the average alpha-band amplitude values of the ipsilateral and contralateral hemispheres in each task. This analysis revealed significant differences in alpha activity between the two hemispheres in the endogenous attention task from 650 – 900 ms (all *p*s < 0.04), with marginally non-significant alpha activity stretching from 900 – 1000 ms (all *p*s < 0.09). However, significant lateralized alpha activity was present much earlier in the hybrid attention task, lasting from 150 – 1000 ms (all *p*s < 0.04), replicating Exp. 1.

Second, in order to compare the magnitude of this lateralized alpha activity across tasks, pairwise comparisons were performed on the alpha amplitude difference values (contralateral minus ipsilateral alpha amplitude) of each task in the a priori defined time windows of the ACOP (260 – 360 ms) and LDAP (500 – 800 ms). These comparisons revealed that the amplitude of lateralized alpha activity was higher in the hybrid than endogenous attention task at the early time window, *t*(15) = 2.69, *p* = 0.02, *d* = 0.67; this difference remained present numerically at the later time window (500-800 ms), but was marginally non-significant then, *t*(15) = 1.92, *p* = 0.07, *d* = 0.48. In sum, these results reveal the presence of lateralized alpha activity following informative cues, with this activity emerging more quickly and with greater magnitude following peripheral vs. central informative cues.

Overall, these data are consistent with the hypothesis that lateralized changes in alpha activity are a general neural marker of visual-cortical enhancement following a cue, regardless of cue format or informativity. The two experiments indicate that changes in occipital alpha rhythm are sensitive to both the time course of attentional deployment following a cue and the spatial information carried by that cue.

## Discussion

A classic distinction in the attention literature is that between endogenous, or voluntary attention, and exogenous, involuntary attention. The differentiation of these two attention systems is well-motivated, as they are each initiated by different events, differ in terms of their temporal dynamics, and are implemented in separate (though partially overlapping) brain networks (Kröse & Julesz, 1989; Müller & Rabbitt, 1989; Nakayama & Mackeben, 1989; Cheal & Lyon, 1991; Corbetta & Shulman, 2002; Peelen, Heslenfeld, Theeuwes, 2004; Chica et al., 2013). However, despite their differences, they each result in similar behavioral effects – improving perception of stimuli appearing at the attended location relative to unattended locations (for a review, see Carrasco, 2011). Here, we investigated the neural processes underlying each type of attention, and asked how they interact to jointly influence behavior. We ran two within-subject experiments that systematically controlled for differences in task design between endogenous and exogenous attentional cueing paradigms by varying only the two main dimensions over which endogenous and exogenous attention tasks typically differ: cue informativity and cue format. By introducing a novel hybrid attentional cue (peripheral, informative cues), we were able to tease apart neural activity related to the shifting of spatial attention and the anticipatory biasing of visual-cortical activity for each type of attention, and also assess their interaction.

Our data reveal that endogenous and exogenous attention exert similar influences over parietal and occipital cortices in response to a cue and prior to the onset of a target. First, oscillatory alpha activity was decreased over occipital cortex contralateral vs. ipsilateral to the attended side following all of the cues. These changes showed a similar contralateral, parietal-occipital focus across the tasks, consistent with the hypothesis that they represent the same visual-cortical enhancement in preparation for a potential target (i.e., a baseline shift in cortical excitability). Secondly, we observed positive deflections over parietal-occipital cortex in the ERP waveforms following informative and uninformative peripheral cues (the ACOP) and informative central cues (the LDAP). These slow-wave ERPs differed in terms of spatial topography and magnitude, suggesting that they represent partially distinct processes. Both lateralized alpha and slow-wave ERPs showed different temporal dynamics following each cue type, providing important insights into their functional roles during the deployment of spatial attention. Specifically, our data are consistent with the slow-wave ERPs reflecting the spatial orienting response of attention and lateralized alpha activity representing the spatially selective biasing of visual cortex activity that is sensitive to, but does not depend on, endogenous topdown signals.

Several studies have shown that alpha activity decreases over contralateral occipital cortex with respect to a voluntarily attended location following a central attention cue (Worden et al., 2000; Rihs, Michel, & Thut, 2007; Jensen & Mazaheri, 2010; Doesburg, Bedo, & Ward, 2016). The observed changes in alpha activity have been interpreted as reflecting top-down anticipatory visual-spatial attention signals that prepare visual cortex to bias subsequent inputs in favor of the attended location – and have often been interpreted as an important index of endogenous attention in particular (Klimesch et al., 1998; Worden et al., 2000; Thut et al. 2006; Doesburg et al., 2016). However, recent studies have demonstrated that peripheral, salient sounds can elicit quick but relatively transient lateralized changes in alpha frequency activity, from ~200 to 400 ms after a cue in the absence of any visual information (Störmer et al, 2016; Feng et al., 2017; see also, Bacigalupo, & Luck, 2019). Still, to this point it has been unclear how lateralized alpha observed during endogenous spatial attention tasks relates to the alpha changes triggered by salient peripheral cues. The present results indicate that lateralized alpha can be triggered rapidly by peripheral cues, regardless of their informativity, but sustains throughout the cue-target interval only when a cue carries spatial information about the target. In particular, we found that lateralized alpha activity was already present at about 150ms after the peripheral cues, but that it emerged later for symbolic cues (at about 650ms) and persisted throughout the entire cue-target interval only when the cue was predictive of the subsequent target location. Importantly, both early and late lateralized alpha changes were focused over parietal-occipital areas, consistent with the hypothesis that they index a common process of enhancing activity in visual cortex even when triggered by distinct cues. This spatially selective pre-activation of visual cortex in anticipation of an impending target stimulus is one of the main principles of the biased-competition model of attention (Desimone & Duncan, 1995), effectively facilitating processing of attended items by biasing neural processing even before any stimulus is presented (Kastner et al., 1999). The present data show that such anticipatory biasing occurs across exogenous and endogenous attention tasks and is possibly implemented by the same mechanism indexed by the occipital alpha rhythm. Thus, these data also add to our understanding of occipital alpha oscillations. First, lateralized alpha over parietal-occipital cortex appears to track the location and time course of both endogenous and exogenous spatial attention. Second, the rapidly emerging portion of alpha – triggered by peripheral cues – is independent of top-down goals, while the later sustained portion is modulated by current task goals.

Of particular interest was how these changes in alpha activity interact with each other when both exogenous and endogenous attention are engaged. Our novel hybrid attention task shows that the biasing activity indexed by alpha oscillations can be effectively “handed off” from exogenous to endogenous attention when an attention-grabbing cue also contains task-relevant information. As can be seen in the hybrid task plots of Figure 2, there is no evidence of a discontinuity in alpha activity following the informative, peripheral cues. Additionally, the topography of this alpha activity remains constant. In other words, it appears that the biasing activity engaged by each system can be effortlessly coordinated when both types of attention are engaged by a novel cue that combines aspects of exogenous and endogenous cueing paradigms.

It is worth noting that this hybrid attention task not only allowed us to disentangle the influence of cue format and informativity on lateralized alpha activity, but also represents a more ecologically valid cueing paradigm. In everyday life, salient events are often predictive of objects that we want to pay attention to. As such, it seems particularly adaptive for exogenous spatial attention, which may initially be captured by a salient event, to exert the same influences on visual-cortical processing as later-arriving effects of endogenous attention in order to optimize stimulus selection. Indeed, the cue-elicited alpha activity of the hybrid attention task indicates that endogenous and exogenous attention work together to orient and sustain attentional biasing at a location. This argues strongly against fully segregated systems underlying each type of attention and demonstrates a novel example of how these two systems may coordinate in the real world.

The present results also illuminate the functional significance of the positive slow-wave ERPs typically observed in response to attentional cues, plotted in Figure 3. Peripheral cues elicited an early positive deflection from ~250 – 550 ms over parietal-occipital cortex. This deflection was focused contralateral to the cue location and was independent of the cue’s spatial informativity (i.e., ACOP). Analogously, a smaller and later lateralized parietal positivity was observed from ~ 400 – 600 ms following the central symbolic cues (i.e., LDAP). These ERP components have been reported previously, and they have each been linked to processes of exogenous and endogenous attention respectively (McDonald et al., 2013; Van Velzen, Forster, & Eimer, 2002). Though both of the ERP components appeared as lateralized positive deflections in the ERP waveform, they varied substantially in timing and magnitude as well as their topographical distributions. Therefore, while it has previously been proposed that both of these components may reflect the same attentional process, simply shifted in time (Hillyard et al., 2016), the current data suggest that this is not necessarily the case.

In the present study, we found the ACOP to be distributed across both parietal and occipital scalp sites, possibly indicating that it reflects a combination of the orienting response and initial biasing in visual cortex (McDonald et al., 2013; Feng et al., 2014; Hillyard et al., 2016). Our results extend previous findings by demonstrating that the ACOP occurs regardless of the top-down significance of a cue. Specifically, in the hybrid attention task, where the peripheral cues were both salient and informative, the ACOP did not differ in magnitude or time course from the positivity elicited by the uninformative peripheral cues of the exogenous task. The similarity of the ACOP following both informative and uninformative peripheral cues indicates that the ACOP is robust to changes in endogenous attention, offering a novel example of the reflexive nature of exogenous attention.

While the ACOP was distributed across both parietal and occipital areas, the LDAP was evident only over parietal areas. The parietal focus of the LDAP in the present study, together with the finding that it dissipates prior to the onset of the target, is consistent with an account of the LDAP as indexing the orienting of attention to a symbolically cued location (Nobre, Sebestyen, & Miniussi, 2000; Van Velzen, et al., 2002; Green & McDonald, 2006). Indeed, the absence of visual-cortical activation during the LDAP window further rules out an account of the LDAP as reflecting the anticipatory biasing of visual processing (Hopf & Mangun, 2000; Kelly et al., 2010). This interpretation explains why the LDAP was absent when a peripheral and spatially informative cue was presented in the hybrid tasks of Experiment 1 and 2: attention was already oriented to the peripheral location by the time the LDAP is usually observed. Critically, the sensitivity of the LDAP to whether attention was already deployed exogenously to a location is a further demonstration of how endogenous and exogenous attention cooperate (and sometimes compete) to determine how perception is influenced following a cue. Therefore, these findings demonstrate that the ACOP and LDAP reflect a shared process of attention – the initial orienting response – but that the occipital activation observed in the ACOP may uniquely represent early and reflexive biasing of neural activity in visual cortex by exogenous attention.

Across our experiments, both lateralized ERPs and lateralized alpha closely followed the purported time courses of each type of attention. There was a rapid orienting response and transient visual-cortical enhancement following exogenous attentional cues; and there was a relatively later orienting response and longer-lasting visual-cortical enhancement following endogenous attentional cues. This nicely demonstrates the temporal sensitivities of these neural markers across different spatial attention tasks. However, both the ERPs and later alpha activity (starting ~450 ms) also differed significantly in amplitude in each of our tasks. While it is possible that these amplitude differences represent a true divergence between the magnitude of neural activity elicited by endogenous and exogenous attentional cues, it also may be the case that differences in the trial-by-trial temporal dynamics of endogenous vs. exogenous attention explain these differences. Specifically, previous research suggests that the endogenous orienting of attention consists of several additional processes relative to the exogenous orienting of attention: such as interpreting the symbolic cue, mapping it to the corresponding target location, and planning the shift of attention to the appropriate location (Hazlett & Woldorff, 2004). The exact timing of these interpretation-and-mapping processes likely varies across trials, and this temporal variability could underlie the differences in magnitude of the peripheral and symbolic cueing effects observed here. Presumably, there is much less temporal variation accompanying the exogenous orienting of attention – where no additional mapping or planning processes are required. Thus, at this point it is difficult to disambiguate whether the differences in magnitude of ACOP and LDAP and lateralized changes in alpha activity are due to actual differences in the size of the effects, or whether they are simply a result of larger trial-by-trial variability in attentional shift time for endogenous relative to exogenous attention. However, future work manipulating the interpretability of a cue as well as the difficulty of the cue-location mapping process may help to distinguish between these accounts.

Overall, our data demonstrate that the orienting of spatial attention triggers lateralized changes in occipital alpha activity and slow deflections in the ERP waveforms – regardless of whether attention is shifted endogenously or exogenously. By directly comparing exogenous and endogenous attention separately and jointly (using a hybrid task), we were able to show distinct functional roles of the lateralized ERPs, which appear to represent the spatial orienting response, and lateralized alpha, which reflects the selective enhancement of visual-cortical activity in preparation of an impending target. The finding that lateralized alpha activity emerges following different cue types, albeit at different time scales, suggests that endogenous and exogenous attention are – at least in part – supported by the same anticipatory visual-cortical biasing mechanisms, enabling them to effortlessly work together to promote most effective stimulus processing.

